# Probing Neuro-Endocrine Interactions Through Wireless Magnetothermal Stimulation of Peripheral Organs

**DOI:** 10.1101/2021.06.24.449506

**Authors:** Lisa Y. Maeng, Dekel Rosenfeld, Gregory J. Simandl, Florian Koehler, Alexander W. Senko, Junsang Moon, Georgios Varnavides, Maria F. Murillo, Adriano E. Reimer, Aaron Wald, Polina Anikeeva, Alik S. Widge

**Author notes:** **Corresponding authors:**, **Dr. Polina Anikeeva**, 77 Massachusetts Avenue, Room 36-812, Cambridge, MA 02139, **Dr. Alik S. Widge**, 2001 6^th^ St SE, Room 3-208, Minneapolis, MN 55455.

## Abstract

Exposure to stress alters hypothalamic-pituitary-adrenal (HPA) axis reactivity; however, it is unclear exactly how or where within the HPA pathway these changes occur. Dissecting these mechanisms requires tools to reliably probe HPA function, particularly the adrenal component, with temporal precision. We previously demonstrated magnetic nanoparticle (MNP) technology to remotely trigger adrenal hormone release by activating thermally sensitive ion channels. Here, we applied adrenal magnetothermal stimulation to probe stress-induced HPA axis changes. MNP and control nanoparticles were injected into the adrenal glands of outbred rats subjected to a tone-shock conditioning/extinction/recall paradigm. We measured MNP-triggered adrenal release before and after conditioning through physiologic (heart rate) and serum (epinephrine, corticosterone) markers. Aversive conditioning altered adrenal function, reducing corticosterone and blunting heart rate increases post-conditioning. MNP-based organ stimulation provides a novel approach to probing the function of HPA and other neuro-endocrine axes and could help elucidate changes across stress and disease models.

## Introduction

Multiple psychiatric disorders, particularly stress and trauma-related conditions, are associated with dysregulation in the hypothalamic-pituitary-adrenal (HPA) axis (1–4), which mediates homeostatic stress responses. Systemically-released adrenal stress hormones, cortisol (CORT; or corticosterone in rodents) and epinephrine (E), are key elements in this pathway. Once the HPA axis is activated, homeostasis is maintained through negative feedback to prevent overstimulation. Circulating adrenal hormones reduce central secretion of their triggers such as corticotropin-releasing hormone (CRH) or adrenocorticotropic hormone (ACTH). Disruption of this homeostatic loop, often by prolonged stress, is commonly found in depression and post-traumatic stress disorder (PTSD)(1,5–7). The consequences of chronic stress exposure or HPA axis dysregulation are mixed, with some studies reporting hyper-responsivity (increased CORT release), and others reporting hypo-responsivity, or a blunted CORT response (3,8–10). These inconsistencies may reflect individual differences in stress sensitivity, and they highlight the need for a more nuanced understanding of HPA function under stress. Specifically, methods are needed for probing the capacity for adrenal release and feedback adaptation at multiple points along the stress and recovery trajectory.

Circulating adrenal hormones also affect learning. Memories formed during states of high emotional arousal can persist and be reactivated more efficiently than memories formed during low arousal (11). These memory-enhancing effects can be adaptive or problematic, depending on the specific memory and its degree of generalization. For instance, trauma-related disorders involve formation of extinction-resistant emotional memories that then lead to pathological avoidance habits (12). There is great interest in finding ways to alter or augment extinction learning processes that could oppose these traumatic memories (13). Pre-clinical studies have suggested timed brain stimulation (14–17), glutamatergic agonists (18,19), and timed increases in adrenal hormones (1–5) as potential strategies for augmenting extinction. Invasive brain stimulation presents a challenge to routine clinical practice, and pharmacologic strategies have shown limited effectiveness in formal trials (20–22). Part of the challenge is that manipulations may need to precisely coincide with the formation of extinction memories (16). This is the basis of a recently-approved brain stimulation treatment for obsessive-compulsive disorder (23). Methods for precisely timed adrenal release would enable pre-clinical studies of HPA effects on extinction learning.

We recently demonstrated an approach for temporally precise adrenal hormone control: direct, wireless adrenal gland stimulation (24). Biocompatible, non-toxic magnetic nanoparticles (MNPs) composed of iron oxide can be injected into adrenal glands (or almost any peripheral organ). In humans or larger animal models, that injection could be performed under X-ray, magnetic resonance, or ultrasound guidance, i.e. as a minimally invasive procedure. In the presence of alternating magnetic fields (AMF), the MNPs dissipate heat. This, in turn, opens native thermosensitive calcium-permeable ion channels from the transient receptor potential (TRP) family, depolarizing electrogenic cells (Figure 1A). This approach increases circulating epinephrine and corticosterone in rats (24), controlled by calcium influx into adrenal cells. This technology offers advantages over other means of probing peripheral organ function. It permits access to tissues where chronic indwelling hardware (e.g. electrodes, catheters) may be difficult to implant or secure. It can be applied without a tether, implying that it could be used in multi-animal assays such as social interaction. Magnetic fields readily penetrate deep into tissue, and using magnetic nanomaterials as transducers enables spatially restricted stimulation. This contrasts with inductive and ultrasonic approaches, where resolution and penetration depth are inversely correlated (25–29). Magnetic activation has advantages over other forms of hardware-free control such as chemogenetics, as it permits tight temporal control over organ stimulation (30).

**Figure 1.**
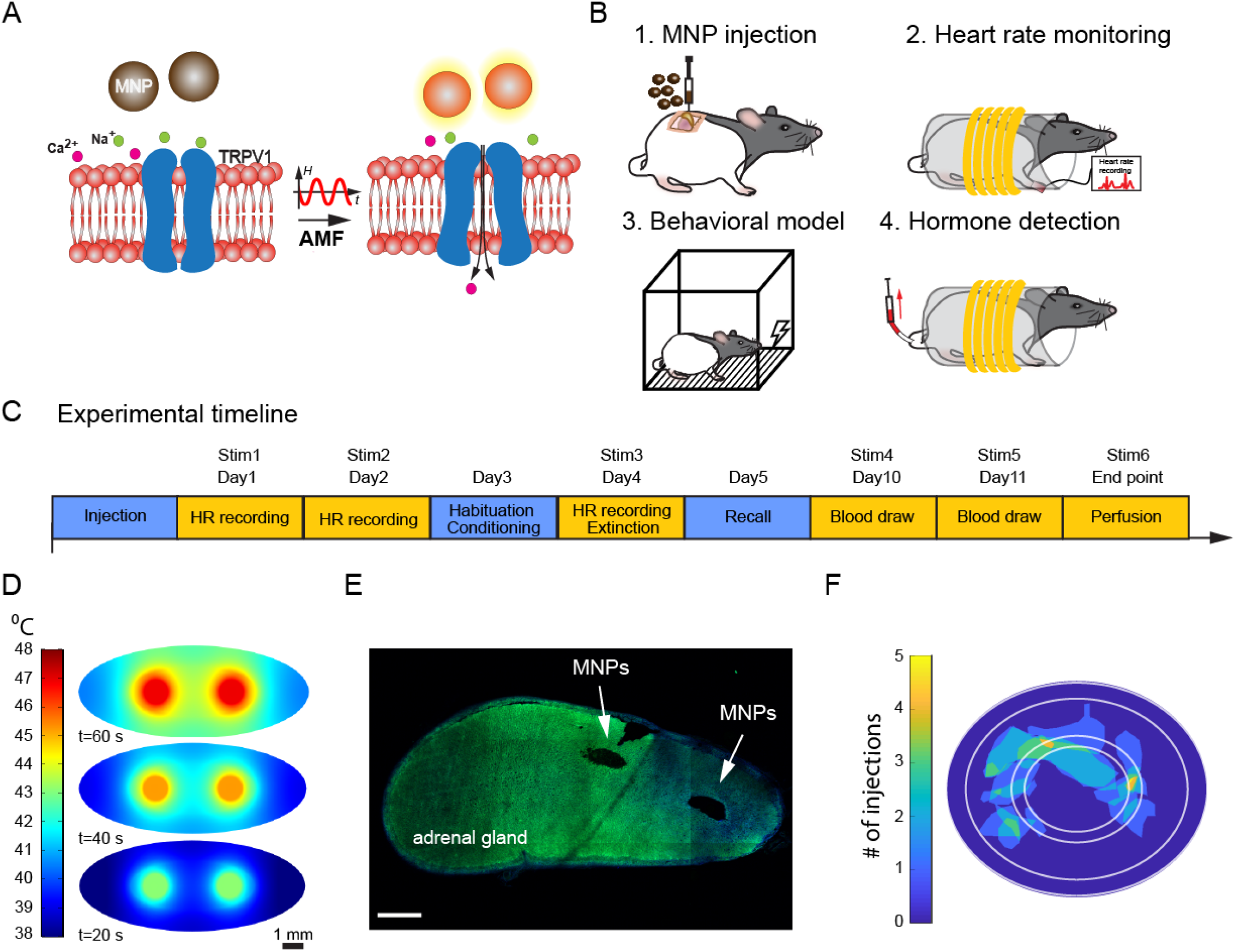
Experimental design and adrenal histology. **(A)** When an alternating magnetic field (AMF) is applied, the magnetic nanoparticles dissipate heat that causes the opening of heat-sensitive calcium ion channels (TRPV1 receptors). The calcium influx causes E and CORT release from adrenal cells. **(B)** Schematic diagram of the procedures that were performed in order. 1. Adrenal MNP injections. 2. Heart rate monitoring before and during magnetothermal stimulation. 3. Aversive conditioning and extinction. 4. Serum collections during magnetothermal stimulation for hormone analyses. **(C)** Experimental timeline. Following adrenal MNP injections, heart rate monitoring took place during MNP stimulation for 2 days (days 1 and 2) before the 3-day behavioral testing (days 3-5). Five days later, blood was collected for serological hormone (E and CORT) measurements (days 10 and 11). **(D)** Finite element modeling of temperature increases at 2 small-volume MNP injection sites within the adrenal gland at 20-s, 40-s, and 60-s of AMF applied. Scale bar = 1mm. **(E)** Representative mosaic of an adrenal gland section with two MNP injections (two black holes). Scale bar = 500 μm. **(F)** A map of MNP and WNP injection sites in the adrenal gland across all animals (n=16).

In this study, we demonstrate the use of that technology to probe HPA axis function over time. We show that an acute stressor (threat conditioning) alters HPA axis function and provide pilot evidence that related behavior (extinction) can be modulated by timed adrenal hormone release.

## Methods and Materials

### Nanoparticle synthesis and characterization

Iron oxide nanoparticles (NPs) were synthesized according to a previously published protocol. We synthesized wüstite (control NPs) or magnetite NPs (active MNPs) from sodium oleate and FeCl_3_∙6H_2_O in a multi-step heating, drying, and solvent control process. All NPs were redispersed in 5 ml of chloroform followed by surface functionalization using polyethylene glycol coating to make NPs biocompatible (see Supplement). Control NPs have the same physical dimensions and chemical elements as MNPs, but are non-magnetic (weakly antiferromagnetic) and do not exhibit hysteretic heating in AMFs (24,31).

### Subjects

We tested magnetothermal stimulation in male Long Evans rats weighing 250-300g. We did not plan a fully-powered study for behavioral effects, but estimated the number of animals per group based on expected attrition (see Supplement). Rats were maintained on a 12 h light/dark cycle and ad libitum chow and water. Experiments were performed during the light phase of the light/dark cycle. All procedures were approved by the Subcommittee on Research Animal Care at the Massachusetts General Hospital (an Institutional Animal Care and Use Committee).

### Adrenal MNP injections

The left adrenal gland of all rats was injected with MNPs or control NPs, under isoflurane anesthesia, after direct visualization and immobilization of the gland. We used a unilateral injection to reduce surgical burden. 1 μl of nanoparticle solution (40 mg_[Fe]_/ml) was injected at a rate of 0.1 μl/min in at least two different locations, for a total volume of 2 μl. Rats recovered from surgery for at least 7 days before entering the experiment. To define the volume and concentration needed for the *in vivo* injections, we created a finite element model to account for the heat distribution within the gland considering the gland dimensions, blood perfusion and a fat layer surrounding the gland. The model was developed using Pennes’ bio-heat equation and similar parameters as in Rosenfeld et al. (2020). We modified the previous model to fit the current experiment (see the Supplement for the table with values).

### Magnetothermal stimulation

We measured HPA axis function by acute magnetothermal stimulation of awake, restrained rats (Figure 1B). Rats were placed inside a plastic restraint tube, to which they had been habituated. The tube was then placed inside the bore of an *in vivo* alternating magnetic field (AMF) stimulation coil, which was a replica of that used in Rosenfeld et al. (2020). The coil was driven by a custom resonant circuit and was actively water cooled (24). At an AMF frequency of 623 kHz, the AMF amplitude values were matched to reach specific loss power (SLP, heating efficiency) of at least 600 W/g in MNPs as in the prior study (24). For all stimulation procedures, rats received 1 minute of active stimulation. Stimulation parameters were selected based on the amount of heat and its spread in the adrenal gland (Figure 1D). In addition, these animals were larger/older than those in Rosenfeld et al. (2020); therefore, we increased the stimulation time from the 40 seconds used in that prior paper, following the same heating model (Figure 1D).

### Heart rate monitoring

To verify that adrenal hormone (particularly epinephrine) release occurred in response to MNP heating, we measured stimulation-induced heart rate changes. A pulse oximetry sensor was attached to the paw after each animal was restrained, and heart rate (bpm) was recorded before, during, and after stimulation using a PhysioSuite (Kent Scientific, Torrington, CT). The experimenter performing stimulation and heart rate procedures (MFM) was blind to each rat’s experimental condition (MNP vs. control NP).

### Behavior

To assess the effects of adrenal stimulation on emotionally valenced learning, we employed a tone-shock conditioning/extinction/recall paradigm that we (32,33) and others (16,34) have previously used to test putative learning and memory enhancers. The experimenter (MFM) was blind to each rat’s experimental condition. Similar to prior experiments, behavior included habituation to the apparatus, tone-shock conditioning, tone-only extinction, and tone-only extinction recall (Figure 1C). Immediately prior to extinction, each animal received 1 minute of magnetothermal stimulation within the coil described above. Pre- and post-stimulation blood samples were collected 6 and 7 days after extinction. We quantified defensive behavior (freezing) by video analysis (ANY-maze, Stoelting Co., Wood Dale, IL).

### Serum hormone quantification

Five days after behavioral testing, we further verified hormone release by lateral tail vein blood collections before and immediately after stimulation (Figure 1C). We previously verified stimulation-induced hormone release properties in unconditioned animals in Rosenfeld et al. (2020). As such, we collected blood samples only after the conditioning and extinction sessions in the current study. Serum hormone levels were quantified via ELISA (MyBioSource, Epinephrine: MBS264776, Corticosterone: MBS761865).

### Histology and image analysis of adrenal glands

Following the experiment, rats were sacrificed via injections of pentobarbital/phenytoin at 150 mg/kg pentobarbital (Beuthanasia®-D C IIIN, Merck Animal Health, Patterson Veterinary, Devens, MA) and transcardial perfusion with 4% paraformaldehyde. Adrenal glands were sectioned in agarose with a thickness of 40 μm and mounted on glass slides to be imaged in a laser scanning confocal microscope (Fluoview FV1000, Olympus). The percentage of nanoparticle coverage of each adrenal sub-region was determined from mosaic scans of the entire adrenal slice via image analysis as in Rosenfeld et al. (2020). Post-processing transformed each adrenal slice to an ellipse with the same semi-axes. This permitted labeling of adrenal sub-regions: medulla, zona glomerulosa (ZG), zona fasciculata (ZC) and zona reticularis (ZR). A map of injection locations was generated across 16 adrenal glands injected with NPs and defined the percentage of area covered with MNPs for each injected gland (Figure 1E, F).

### Statistical analysis

All analyses were performed using RStudio. As noted above, this study was not powered to detect significant between-group differences, and we present parametric statistics primarily to demonstrate possible effect sizes. See the Supplement for the rationale for each analytic choice.

We analyzed heart rate for Days 1 & 2 (pre-conditioning) and Day 4 (post-conditioning) separately, retaining only animals with adequate data and verified NP placement. We summed samples during stimulation (300 to 360 sec) to compute the area under the heart rate curve (AUC). We compared AUC of active and control conditions using a generalized linear model with gamma distribution, identity link function, and a single independent variable (treatment condition).

After excluding animals for experimental failures and outlier values, we analyzed the mean of the serum hormone levels across the two days of collection. We converted the data to a post-stim/pre-stim ratio, which we compared between active and control animals with a two-sample t-test on the log of the post/pre ratio.

We quantified freezing behavior as the percentage of the 30s CS tone that was spent freezing. Data were normalized to each rat’s individual baseline and smoothed using a centered moving average. Freezing was then analyzed in a beta regression using trial, testing phase, and treatment (fixed effects, including 2-way and 3-way interaction terms) as explanatory variables. As an additional unplanned analysis, we tested whether active vs. control animals differed in their freezing at the end of extinction (Trials 18-20), using an independent-samples t-test.

## Results

### MNP influences on heart rate

Prior to tone-shock conditioning, animals injected with active MNPs showed adrenal engagement in response to an AMF stimulus, evidenced by an increased heart rate during stimulation compared to controls (Figure 2A). The between-group difference did not reach significance in this small sample [*t*(14)=0.807, *p*=0.433]. Further, aversive conditioning changed adrenal responsivity. After conditioning (but before extinction on day 4), the same stimulation produced no visible difference in heart rate between the control and active MNP animals [*t*(14)=− 0.025, *p*=0.98] (Figure 2B). Heart rate rose more slowly in the animals injected with active MNPs, and there was greater variability as stimulation continued (increasing width of error bars specifically during the stimulation period).

**Figure 2.**
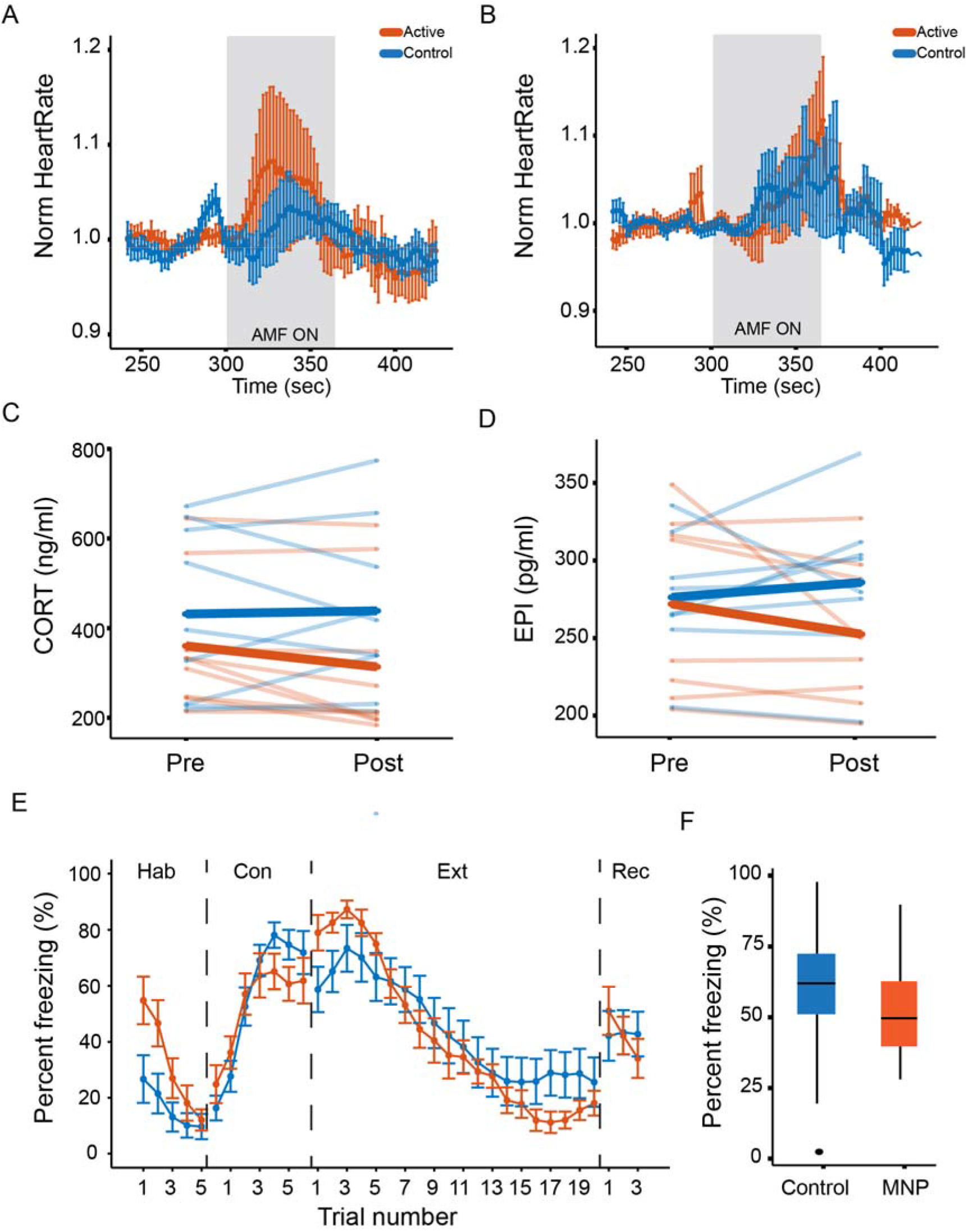
Magnetothermal adrenal stimulation effects on heart rate, hormone levels, and freezing behavior. **(A, B)** Heart rate measurement during magnetothermal stimulation on (A) days 1and 2 before conditioning and (B) on day 4 after conditioning. **(C, D)** Serological analysis. (C) Corticosterone and (D) epinephrine were measured before (pre) and after (post) magnetothermal stimulation. **(E)** Percent freezing per trial across the habituation, conditioning, extinction, and recall phases of the behavioral paradigm (controls, n=13; active, n=10). **(F)** Percent freezing averaged across the last three trials of extinction (Trials 18-20) for control and active MNP rats.

### MNP modulation of adrenal hormones

After aversive conditioning and extinction, the circulating hormone response to magnetothermal stimulation was also altered. In animals injected with active MNPs, serum CORT and E both decreased from pre- to post-stimulation (Figure 2C, D). The change in CORT reached statistical significance compared to control MNPs [*t*(16)=−2.2, *p*=0.047], whereas the change in E did not [*t*(14)=−1.8, *p*=0.091], although significance should be evaluated with caution given the sample size. Importantly, these are the reverse of the change we previously reported in unconditioned animals (24).

### MNP modulation of defensive behavior

Freezing to conditioned tones did not significantly differ between the animals injected with active MNPs and those injected with control non-magnetic NPs in any experimental phase (Figure 2E, F). There was a non-significant enhancement of extinction learning from adrenal stimulation at the end of extinction [*t*(20.4)=−0.579, *p*= 0.569], that did not persist to the recall phase.

## Discussion

We previously demonstrated remote control of adrenal hormone release via magnetically triggered heating of locally injected MNPs (24). Here, we demonstrated the use of this technology to probe the state dependence of HPA axis function. After aversive conditioning, the same adrenal stimulation produced visibly different changes in heart rate. This may represent exhaustion of a readily releasable pool. After conditioning, animals may have continuously high adrenergic tone, with little reserve E available for release in response to subsequent stimulation. This may offer a model for persistent hyper-arousal in trauma/stress-related disorders and demonstrates the potential role of this technology as a probe of neuro-endocrine interactions. MNP stimulation also decreased serum E and CORT. In our prior study of non-conditioned animals, the same stimulation increased these hormones at the same time point (24). This aligns with the heart rate results, in that it again represents a blunted response. These decreases potentially reflect an up-regulation of E/CORT metabolism or feedback inhibition as a result of a repeatedly activated HPA axis during conditioning, which could lead to a net decrease by the post-stimulation measurement (35). These results further demonstrate that magnetothermally-driven adrenal release can probe learning-related alterations of HPA axis function.

Behaviorally, we observed only a small difference in freezing, even though circulating adrenal mediators might be expected to improve extinction learning (36–38). The reduction in pre- to post-stimulation serum levels suggests that the lack of a behavioral difference may be driven by a lack of sustained change in circulating mediators, or even by their suppression. The timing of activation/secretion may also be important. Previous studies suggest time-dependent memory enhancement, e.g. hydrocortisone reduced defensive behavior only if given close to the time of training (39). Because our technology allows more precise control over hormone release timing compared to commonly used approaches such as a systemic injection or delivery via an osmotic pump, it could permit dissection of this time dependence.

The current approach was limited to adrenal gland activation by triggering heat-sensitive TRP channels, in a relatively non-specific way across the entire organ, in restrained animals. However, the resulting limitations can be overcome. For example, in tissue without thermosensitive channels, we have activated mechanosensitive ion channels by altering the MNP structure and magnetic field conditions (40). Moreover, magnetothermal stimulation can also inhibit cell activity by activating heat-sensitive hyperpolarizing channels (41). Controlled injections of MNPs to specific substructures within the adrenal gland, instead of the current multiple-injection approach, could increase specificity and permit the independent release of individual hormones. Specificity might also be achieved using a multiplexing approach for controlling different populations of MNPs within the same organ (42). While we used restrained animals, the current coil design would permit experiments with freely moving mice (43). Freely moving rat experiments are possible albeit demand scaling up of the coils and driving power electronics (44).

This technology could generalize beyond the HPA axis. Sex hormones such as estrogen, progesterone, and testosterone all affect learning processes (32,33,45–47) and are broadly implicated in psychiatrically relevant functions (6,48,50). Systemic administration of these hormones in animal studies relies on subcutaneous or intraperitoneal injections, which can be stressful and interfere with the sex hormone effects being investigated. TRP receptors are present in the ovaries, testes, and hypothalamus (49,51,52). Therefore, the hypothalamic-pituitary-gonadal (HPG) axis could be similarly probed by magnetothermal stimulation.

Beyond their use as probes of peripheral organ function, magnetic nanomaterials have been increasingly recognized as therapies and “theranostic” tools. They are being applied for treating cancer, diagnosing atherosclerosis, delivering drugs, etc. (53,54). However, their utility as tools for neuropsychiatric treatment and investigation is less defined. While intracranial or deep brain MNP stimulation has mostly been validated with motor behaviors (55), these particles may be a valuable tool for creating and/or manipulating models of neuropsychiatric disorders (43,56). For instance, MNP stimulation of the prelimbic cortex reduced immobility in the forced-swim test and increased sucrose consumption in stressed mice (56). Rao et al. (2019) demonstrated the use of MNPs for targeted drug delivery, where MNP-stimulated drug release increased dopamine-mediated social behavior. While these findings illustrate how MNP stimulation can be used to alter behavior via central modulation, the present study suggests a potential for modulation in the periphery, which may be more accessible and translatable.

Magnetothermal or magnetomechanical approaches are not universally useful. Chemogenetics will be simpler wherever temporal precision is not necessary. Very complex arenas will be difficult to instrument with coils for MNP activation, favoring more mature, tethered technologies such as optogenetics. The coils require specialized driving hardware and high-voltage supply lines that may not be available in all laboratory environments. It would similarly be hard to run many animals at once using this paradigm, and pharmacological/chemogenetic tools would be more suited if high throughput is needed. As nanomaterials, MNPs still need to be placed into organs of interest and may degrade, migrate, or be eliminated over time, unlike larger optogenetic fibers (57,58).

Despite the limitations described above, we have demonstrated the use of MNPs to remotely and more precisely detect and assess alterations within a peripheral stress system. Future research could further examine the timing of MNP peripheral stimulation effects on behavior, influences in the brain, and their potential role in the restoration of post-conditioning adrenal function. As the technology becomes more refined and widely available, these on-demand peripheral release approaches could provide valuable new tools for understanding and eventually altering the biology of mental illness beyond the brain.

## Acknowledgements

This work was funded in part by the DARPA ElectRx Program (HR0011-15-C-0155 under Dr. D. Weber), the Bose Research Grant, and the NIH BRAIN Initiative (1R01MH111872). This work made use of the MIT MRSEC Shared Experimental Facilities under award number DMR-14-19807 from the NSF. D.R. is a recipient of the MIT-Technion Fellowship. A.S.W. was additionally supported by the MnDRIVE Brain Conditions Initiative.

## Disclosures

Dr. Widge, Dr. Anikeeva, and Dr. Rosenfeld are inventors on a patent application for therapeutic uses of controlled adrenal release. Dr. Maeng, Gregory J. Simandl, Florian Koehler, Dr. Senko, Dr. Moon, Georgios Varnavides, Maria F. Murillo, Dr. Reimer, and Aaron Wald do not have any financial or other conflicts of interest to declare.

## citations for the RStudio packages

### betareg

Francisco Cribari-Neto, Achim Zeileis (2010). Beta Regression in R. Journal of Statistical Software 34(2), 1-24. URL http://www.jstatsoft.org/v34/i02/.

### broom

David Robinson, Alex Hayes and Simon Couch (2020). broom: Convert Statistical Objects into Tidy Tibbles. R package version 0.7.3. https://CRAN.R-project.org/package=broom

### fitdistrplus

Marie Laure Delignette-Muller, Christophe Dutang (2015). fitdistrplus: An R Package for Fitting Distributions. Journal of Statistical Software, 64(4), 1-34. URL http://www.jstatsoft.org/v64/i04/.

### ggplot2

H. Wickham. ggplot2: Elegant Graphics for Data Analysis. Springer-Verlag New York, 2016.

### ggpubr

Alboukadel Kassambara (2020). ggpubr: ‘ggplot2’ Based Publication Ready Plots. R package version 0.4.0. https://CRAN.R-project.org/package=gg pubr imputeTS: Steffen Moritz and Thomas Bartz-Beielstein, *The R Journal* (2017) 9:1, pages 207-218. https://cran.r-project.org/web/packages/imputeTS/index.html

### plotrix

Lemon, J. (2006) Plotrix: a package in the red light district of R. R-News, 6(4): 8-12.

### readxl

Hadley Wickham and Jennifer Bryan (2019). readxl: Read Excel Files. R package version 1.3.1. https://CRAN.R-project.org/package=readxl

### tidyverse

Wickham et al., (2019). Welcome to the tidyverse. Journal of Open Source Software, 4(43), 1686, https://doi.org/10.21105/joss.01686

## Notes

https://github.com/tne-lab/magnex.git

