## Supplemental file for "Probing Neuro-Endocrine Interactions Through Wireless Magnetothermal Stimulation of Peripheral Organs"

### Methods and Materials

#### Nanoparticle synthesis.

Iron oxide nanoparticles (NPs), either magnetic (active MNP) or non-magnetic wüstite (control NP) were synthesized by altering the solvent ratios during synthesis. We synthesized wüstite (control NPs) or magnetite NPs (active MNPs) by mixing sodium oleate (95%, TCI America) and 30 mmol of FeCl_3_∙6H_2_O (99%, Acros Organics) were mixed by heating to reflux (60 ºC) in a solvent mixture of hexane, ethanol and ddH_2_O for one hour under N2. The mixture was heated to 110 ºC and dried overnight on a hotplate. MNPs were prepared by degassing the iron-oleate mixture at 90 ºC for 1 h in 2:1 (volume ratio) of 1-octadecene (90%, 10 ml) and dibenzyl ether (98%, 5 ml) and heated to 200 °C under N2. Control NPs were prepared similarly but with 20 ml of 1-octadecene as solvent. Mixtures were then heated to 200 °C under N2 and then to reflux at ~325 °C for 30 minutes. Both types of particles share a size of 22 nm and were transformed by surface functionalization using poly(ethylene glycol) grafted with poly(maleic anhydride-alt-1-octadecene) similar to Rosenfeld et al. (2020). NP concentration was measured using inductively coupled plasma-optical emission spectroscopy ICP-OES (Agilent 5100 DVD) and concentrated to a final concentration at injection of 40 mg _[Fe]_/ml. For estimating the specific loss power (SLP, heating efficiency per gram of iron) of the NPs based on the dynamic hysteresis model (1), we used a similar calorimetry method described previously in our work (2). Briefly, an optical fiber temperature probe (Omega HHTFO-101) was used to measure temperature change in the ferrofluid solution inside a 7.5 mm gap of a toroidal ferromagnetic core that is driven by a custom-built series resonant circuit generating AMF with frequency ƒ = 515 kHz and amplitude H0 = 15 kA/m. The SLP of MNPs in those conditions was estimated by 880 ± 38 W/g[Fe] (mean ± standard deviation, std.) while the same conditions applied to control NPs yield a negligible SLP of 8 ± 4 W/g[Fe] (3).

#### Subjects.

Adult male Long Evans rats weighing 250-300g were obtained from Charles River Laboratory (Wilmington, MA). For this methodological proof of concept, we did not plan a fully-powered study. Rather, we planned for 20 animals per group (active MNP vs. control NP), with the expectation of substantial attrition as we refined the protocol or due to behavioral anomalies (e.g., animals that could not acquire threat conditioning). We estimated based on prior experience that 20 animals per group would ensure at least 10 completers even under attrition, which would permit estimation of effect sizes. Of these 40 initial rats, 28 completed the full protocol and were retained in at least some aspects of the analysis; Figure S1 diagrams the reasons for attrition per analysis.

All rats were housed in the rodent facility at the Massachusetts General Hospital Center for Comparative Medicine in Charlestown, MA. The animals were left undisturbed for 1 week after their arrival to allow for acclimation to the colony room. Following this acclimation period, all animals were handled daily for at least 5 minutes to minimize handling stress. The rats also underwent tube habituation, which consisted of daily exposure to the restraint tubes used during heart rate assessments, blood collections, and magnetothermal stimulation (see below). Sufficient tube acclimation was determined based on heart rate stabilization across training sessions and reduced stress responses, i.e. fewer vocalizations. Rats were maintained on a 12 h light/dark cycle and a diet of ad libitum rat chow and water. Experiments were performed during the light phase of the light/dark cycle. All procedures were approved by the Subcommittee on Research Animal Care at Massachusetts General Hospital and abided by the guidelines set forth by this Institutional Animal Care and Use Committee.

Finite element modeling.

To estimate the needed volume and concentration of MNPs and the number of injection sites for the in vivo experiments that will allow to reach the threshold temperature of TRPV1 ion channel (42 ºC), we created a finite element modeling of heat distribution relying on the Pennes’ bio-heat equation similar to Rosenfeld et al. (2020). The chosen dimensions for the glands were modeled as an ellipsoid of 5.5×2.2×2.2 mm^3^ (with additional fat layer) and the following equations:

ρ_A_C_A_$\frac{\partial T}{\partial t}$=K_A_∇^2^T+ ρ_b_C_b_w_b_(T-T_b_)+P r {adrenal tissue}

Ρ_F_C_F_$\frac{\partial T}{\partial t}$=K_A_∇^2^T+ ρ_b_C_b_w_b_(T-T_b_)+P r {fat tissue}

With the following values (Supplementary table 1):

| **Parameter** | **Value** |
| --- | --- |
| Specific loss power of MNPs | 600 W/g _[Fe]_ |
| MNPs concentration | 40 mg/ml |
| Blood density, ρ_b_ | 1000 kg/m^3^ |
| Heat capacity blood, C_p,b_ | 4180 J/(kg∙K) |
| Blood perfusion rate, ω_b_ | 0.0064 s^-1^ |
| Arterial blood temperature, T_b_ | 37 ºC |
| Initial and boundary temperature, T_o_ | 37 ºC |
| Heat capacity adrenal, C_p,A_ | 3540 J/(Kg∙K) |
| Adrenal density, ρ_A_ | 1020 kg/m^3^ |
| Adrenal thermal conductivity, K_A_ | 0.52 W/(m∙K) |
| Heat capacity fat, C_p,F_ | 2348 J/(Kg∙K) |
| Fat density, ρ_F_ | 911 Kg/m^3^ |
| Fat thermal conductivity, K_F_ | 0.21 W/(m∙K) |

#### Adrenal MNP injections.

All rats were injected with intraperitoneal (i.p.) atropine (0.01 mg/kg) for respiratory support. About 20 minutes after this injection, the rats were placed into an induction chamber and were anesthetized with isoflurane. After induction, animals were injected with subcutaneous (s.c.) narcotic pain reliever buprenorphine (0.05 mg/kg), an anti-inflammatory non-narcotic pain reliever flunixin meglumine (2.5 mg/kg, s.c.), and intramuscular (i.m.) antibiotic enrofloxacin (5 mg/kg). To minimize the amount of surgical burden each animal experienced, only the left adrenal gland was injected with MNPs or control NPs. The incision site was shaved and cleansed with alternating Betadine and ethanol. Once the incision was made, the left adrenal gland was visualized and isolated as in Rosenfeld et al. (2020). The needle of a Hamilton syringe containing the nanoparticles was lowered into the gland. The syringe was connected to a stereotaxic infusion pump and injected with the nanoparticles in a volume of 1 µl and injection rate of 0.1 µl/min in at least two different locations, for a total volume of 2 µl. The needle was kept inside the gland for 10 minutes post-injection to prevent leakage of the ferrofluid solution. Surgeries were performed by LYM, MFM, and DR. LYM and DR were aware of which injections contained control NP vs. MNP, but MFM was blinded. The data analysis was similarly performed blind to condition (see below). Rats recovered from surgery for at least 7 days before entering the experiment.

#### Behavior.

The behavioral testing procedure was run across 3 consecutive days. For each phase, an 82-dB tone conditioned stimulus (CS) was presented. On day 1, all rats underwent a habituation phase that consisted of 5 trials of tone CS-alone presentations. Habituation was immediately followed by the aversive conditioning phase. Conditioning consisted of 7 trials of tone CS and shock unconditioned stimulus (US) pairings. In the conditioning trials, each tone culminated with a 0.5mA footshock administered by the grid floor in the operant chamber. The extinction phase took place on day 2, consisting of 20 tone-CS alone trials. Immediately prior to extinction, each animal received 1 minute of magnetothermal stimulation within the coil described above. Heart rate measurements were taken prior to the start of the experiment. Pre- and post-stimulation blood samples were collected 6 and 7 days after the extinction session.

On day 3, the Recall phase comprised 3 CS-alone trials. During each phase, video recordings were analyzed to assess freezing behavior during the experimental trials. Freezing behavior was analyzed via ANY-maze (Stoelting Co., Wood Dale, IL) post-experimentation using video recordings taken within the operant chamber during each behavioral testing phase (habituation, conditioning, extinction, and recall).

#### Blood and serum collection.

Five days after the end of the behavioral testing, we further verified hormone release by lateral tail vein blood collections before and immediately after stimulation. To increase vasodilation of the lateral tail vein and thus blood flow, tails were warmed for 2 – 4 minutes with hand warmers*.* The animals were then anesthetized with 0.25 mg/kg ketamine to insert a tail vein catheter, then allowed to recover for blood collections prior to and after stimulation. After the catheter was implanted, it was flushed with saline and blocked with heparin (0.2 ml). Blood samples were drawn after removing the heparin block. After at least an hour of sitting at room temperature for the blood to clot, the blood samples were centrifuged at 5,000rpm for 5 minutes at room temperature. We chose to perform this collection only after the conditioning and extinction paradigm because we had already verified pre-conditioning hormone release properties in Rosenfeld et al. (2020). Repeated tail vein collections might also have been highly stressful for the animals (much more so than simply resting in a restraint tube), and thus might have altered the conditioning outcomes.

#### Serum hormone quantification.

Serum was kept in aliquots at -20 °C and thawed on ice prior to the beginning of the ELISA assay. ELISA kits from MyBioSource (Epinephrine: MBS264776, Corticosterone: MBS761865) were used according to the manufacturer protocol. The final quantification was conducted using a plate reader to read the optical density. For each tested hormone, the absolute values were calculated according to a calibration curve, obtained separately for each tested plate.

#### Histology.

At the end of all of the experimental procedures, rats were sacrificed via transcardial perfusion. All rats were intraperitoneally injected with pentobarbital/phenytoin at 150 mg/kg pentobarbital (Beuthanasia®-D C IIIN, Merck Animal Health, Patterson Veterinary, Devens, MA). After confirming that they were sufficiently anesthetized by showing unresponsivity to toe pinch, all rats were transcardially perfused with 0.9% saline (exsanginuation) and 4% paraformaldehyde (fixation) at a rate of about 21 mL/minute. Adrenal glands were collected at the time of perfusions and kept in 4% paraformaldehyde. They were then sectioned with a cryostat to confirm the location of the nanoparticle injections. The percentage of nanoparticle coverage of each area was determined from mosaic scans of the entire adrenal slice and with image transformation analysis as in Rosenfeld et al. (2020).

#### Statistical analysis.

All analyses used RStudio with packages betareg, broom, fitdistrplus, ggplot2, ggpubr, imputeTS, plotrix, readxl, and tidyverse. The complete code for all analyses, with corresponding data files, is available online at <https://github.com/tne-lab/magnex>. For all analyses, we identified appropriate distributions for statistical testing by visualization of the Cullen-Frey graph in fitdistrplus.

##### Statistical analysis – Heart Rate.

We recorded heart rate data before, during, and after stimulation. Due to animal movement and occasional loss of the pulse oximetry signal, heart rate tracings contained gaps. We imputed these missing values using Kalman smoothing (“imputeTS'' package, “na_kalman” function), using the default parameters. Post-imputation, data were scaled by dividing each measurement in the time series by the mean pre-stimulation heart rate. Day 1 and 2 data were averaged at each time point for each respective animal. Day 4 data was left separate because measurements were obtained after the conditioning phase and thus were expected to have had different underlying neurobiology. We retained only animals where we were able to successfully impute missing data. We excluded animals in the active MNP group if the histology showed no particles in the medulla. This left n=11 control and n=9 active rats for this analysis. We summed samples within the stimulation time range (300 to 360 sec) to compute the area under the heart rate curve (AUC). These values best fit a gamma distribution, and thus we compared active and control conditions using a generalized linear model with gamma distribution, identity link function and a single independent variable (treatment condition).

##### Statistical analysis – Serum.

Serum analyses assessed epinephrine and corticosterone levels immediately prior to and after stimulation. We analyzed the mean of the pre-stimulation and post-stimulation samples across the two days of collection. 9 animals were excluded from analysis due to blood sampling issues, e.g. catheter malfunctions. For the remaining animals, the serum values were best described by a beta distribution. We then excluded animals with outlier pre-stimulation hormone levels, defined by fitting a beta distribution to all pre-stimulation hormone values (corticosterone and epinephrine separately). We excluded the serum samples whose baseline was extremely unlikely, i.e. p<0.0005 based on the fitted distribution. This method excluded 1 additional animal. Finally, animals without active MNP placement in the medulla for epinephrine, or the cortex for corticosterone release, were also excluded (animals retained for E, control: n=9, active: n=8; CORT, control: n=9, active: n=9). We then converted the data to a post-stim/pre-stim ratio. These values were consistent with a log-normal distribution for each hormone, hence active and control animals were compared with a two-sample t-test on the log of the post/pre ratio.

##### Statistical analysis – Behavior.

The primary behavioral outcome from the 3-day extinction paradigm was animals’ freezing behavior, expressed as a percentage of the 30s CS tone that was spent in freezing. The extinction paradigm consists of four testing procedures: habituation (day 1), conditioning (day1), extinction (day 2), and recall (day 3). We analyzed freezing with a beta regression model, which is designed for data spanning the 0.00 ≤ x ≤1.00 interval. For each animal, the raw freezing scores were adjusted to span this full range using a min-max normalization ((x - min)/(max-min)). Normalizations were done separately for each phase of the testing procedure. These normalized data were scaled by dividing by each rat’s individual baseline freezing, defined as the average of the final 3 trials of the habituation phase (Trial 3-5). Finally, we smoothed the normalized trial-to-trial data using a centered moving average with window size 3. Post-normalization, we confirmed that freezing data fit a beta distribution. Data then were analyzed in a beta regression using trial, testing phase, and treatment (fixed effects, including 2-way and 3-way interaction terms) as explanatory variables. Before the analysis, we excluded animals that did not initially condition, and thus that could not have experienced extinction learning. Conditioning failures were determined by visual inspection of the conditioning behavior curve, by an investigator (ASW) who was blind to each rat’s treatment assignment. We also excluded animals in the active MNP group where both cortex and medulla were particle-free. These approaches excluded 5 rats (21.7% of total), with n=23 rats retained in the analysis (control: n=13; active: n=10). For the analysis comparing the final three trials of extinction, we confirmed that these averages followed a normal distribution before applying a t-test.

**Figure S1.**

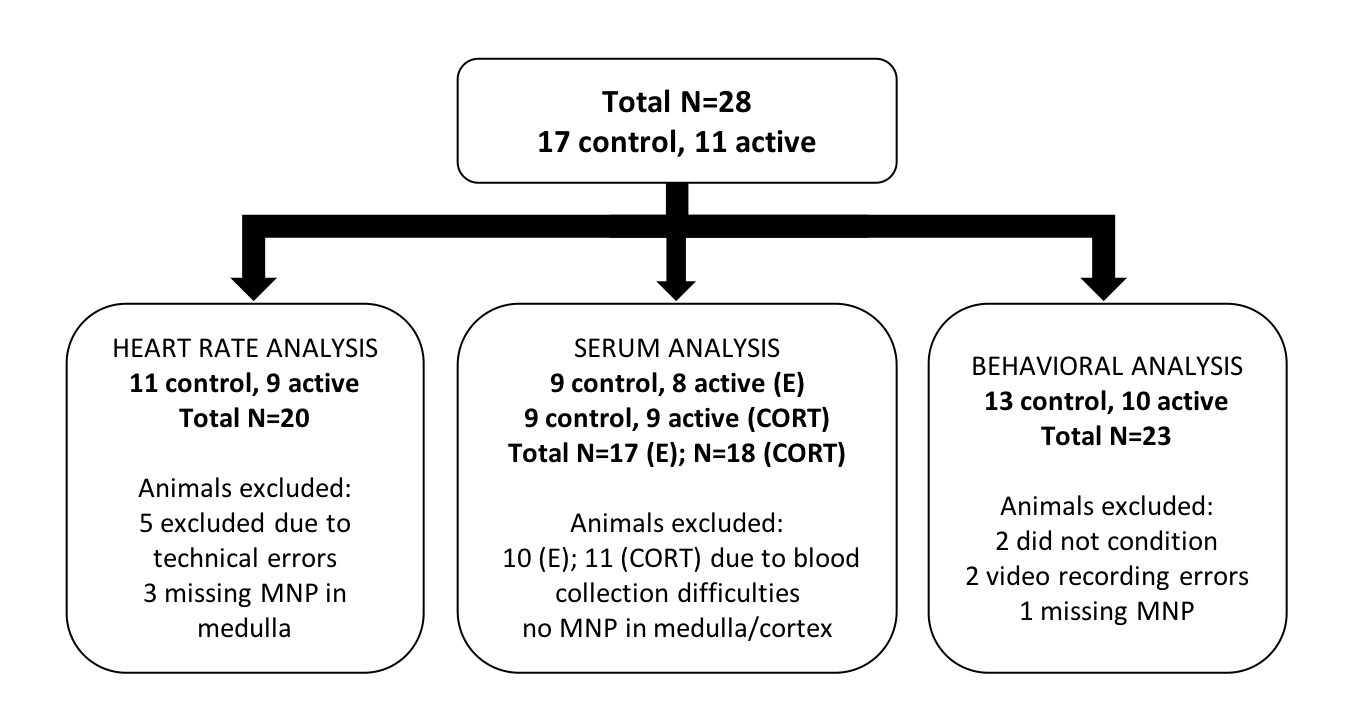

**Figure S1.** Summary chart of exclusions by analysis.
